# Auxin-inducible degradation of UNC-116 in *C. elegans* inhibits bidirectional dense core vesicle transport and worm locomotion on different timescales

**DOI:** 10.1101/2025.06.23.661018

**Authors:** Astrid Boström, Anna Gavrilova, Gino B. Poulin, Victoria J. Allan

## Abstract

The microtubule motor kinesin-1 is vital in neurons, with mutations being associated with neurological diseases. Deletion of the *Caenorhabditis elegans* kinesin-1 gene *unc-116* is lethal, and viable mutants are uncoordinated. Here, we use auxin-mediated degradation to deplete UNC-116 protein at different developmental stages and monitor the effects on cargo transport and locomotion. UNC-116 is substantially degraded within 1 hour of auxin treatment, by which time bidirectional dense core vesicle (DCV) motility is affected. After 4 hours, dynein-driven DCV movement is lost and only limited plus-end-directed DCV motility remains, likely driven by kinesin-3 (UNC-104). DCV movement recovers substantially after rescue from auxin for 24 hours. There is a time-lag between loss of protein and effects on locomotion, as crawling and swimming/thrashing is unaffected until 6-14 hours on auxin. By 18-24 hours, animals are as uncoordinated as the *unc-116(rh24sb79)* mutant. Notably, degradation of UNC-116 in neurons alone inhibits crawling and swimming, revealing the importance of neuronal kinesin-1 for locomotion. Overall, by bypassing early developmental UNC-116 functions, we reveal that UNC-116 is essential for bidirectional DCV transport and crucial for locomotion.

## Introduction

Intracellular transport of cargoes along microtubules is fundamental for maintaining cellular organisation (Cason and Holzbaur, 2022; Sleigh et al., 2019). This process is carried out by two classes of microtubule motors: kinesins and cytoplasmic dynein (dynein). Most kinesins are anterograde motors, transporting cargoes toward the microtubule plus-ends, whereas dynein transports cargoes retrogradely, toward minus-ends (Hirokawa et al., 2009; Reck-Peterson et al., 2018). Intracellular transport is particularly crucial in neurons since components must be transported long distances to reach the length of axons and dendrites (Bentley and Banker, 2016; Cason and Holzbaur, 2022; Guedes-Dias and Holzbaur, 2019).

Kinesin-1, one of the best characterised motors, is responsible for carrying a diverse range of cargoes from membranous cargo to mRNA, and microtubules themselves (Cross and Dodding, 2019; Miki et al., 2005; Verhey and Hammond, 2009). Mammals have three genes encoding the kinesin-1 motor subunit: the ubiquitous *KIF5B* and the neuron-specific *KIF5A* and *KIF5C* (Kanai et al., 2000). Kinesin-1 mutations are associated with disorders including Amyotrophic Lateral Sclerosis, Charcot Marie-Tooth disease, and Hereditary Spastic Paraplegia, leading to eventual spasticity and/or movement difficulties (eg. Baron et al., 2022; Bayrakli et al., 2015; Chiba and Niwa, 2024; Melo et al., 2015; Nam et al., 2018).

*C. elegans* has one isoform of the kinesin-1 motor subunit, UNC-116. Complete loss of UNC-116 function by gene knockout is embryonic lethal, and hypomorphic *unc-116* mutants where UNC-116 protein retains some function are uncoordinated and move very poorly (Gavrilova et al., 2024; Patel et al., 1993; Sakamoto et al., 2005; Yang et al., 2005). This reduced locomotion is not likely to involve defects in synaptic vesicle delivery, as they are transported by the kinesin-3 member, UNC-104 (Hall and Hedgecock, 1991; Kumar et al., 2010; Niwa et al., 2016; Ou et al., 2010; Wu et al., 2013; Zheng et al., 2014). However, kinesin-1 cargoes with a potential role in maintaining locomotion include mitochondria (Chen et al., 2021; Rawson et al., 2014; Sure et al., 2018; Zhao et al., 2021), glutamate receptors (Hoerndli et al., 2013; Hoerndli et al., 2015) and dense core vesicles (Gavrilova et al., 2024). *unc-116* mutant animals also have developmental problems including microtubule polarity reversals in dendrites (Harterink et al., 2018; He et al., 2020) and defects in axonal outgrowth and branching (Aguirre-Chen et al., 2011; Drozd & Quinn, 2023; Gavrilova et al., 2024; Su et al., 2006). The severe locomotion defects seen in *unc-116* mutants could therefore be due to reduced movement of multiple cargoes combined with problems in neuronal development.

Neuronal dense core vesicles (DCVs) deliver and release neuropeptides, neurotrophic factors and other signalling components (Gondré-Lewis et al., 2012). This delivery contributes to many aspects of physiology, including learning, development, locomotion and ageing (Gondré-Lewis et al., 2012; Randi et al., 2023; Ripoll-Sánchez et al., 2023). DCV motility is complex, as they are transported bidirectionally in axons by a combination of kinesin-1, kinesin-3 and dynein (Barkus et al., 2008; Gavrilova et al., 2024; Gumy et al., 2017; Kwinter et al., 2009; Lim et al., 2017). Evidence suggests that transport out of the cell body is carried out by the faster kinesin-3, after which kinesin-1 drives continued transport to the axon tip (Gumy et al., 2017; Lim et al., 2017; Park et al., 2023). Compromising one motor often leads to defective transport in both directions rather than upregulating unidirectional movement, suggesting that antagonistic motors are somehow dependent on each other (Ally et al., 2009; Gavrilova et al., 2024; Hancock, 2014; Kwinter et al., 2009; Lim et al., 2017; Pilling et al., 2006; Yi et al., 2011). The mechanisms governing this “paradox of co-dependence” remain unclear (Hancock, 2014). Investigating the roles of specific microtubule motors in multicellular models is needed to shed light on how bidirectional transport is regulated and enhance understanding of how dysfunction leads to disease phenotypes.

Depleting the UNC-116 protein at specific times during development and once development is complete offers a way to investigate the multi-faceted functions of kinesin-1. To do this, we have made use of the auxin-induced degradation (AID) system, where a protein tagged with a degron amino acid sequence is rapidly degraded upon auxin addition by a plant TIR1 F-box protein that forms an SCF E3 ubiquitin ligase complex with endogenous Skp1 and Cullin (Nishimura et al., 2009). In *C. elegans,* conditional and tissue-specific protein degradation can be mediated when worms expressing TIR1 in specific cells are exposed to auxin (Ashley et al., 2021; Zhang et al., 2015). AID has been used to degrade the dynein motor subunit (Cavin-Meza et al., 2022; Zhang et al., 2015), and the kinesin-3 motor UNC-104 (Cahoon and Libuda, 2021). Very recently, UNC-116 AID has been used to analyse the role of kinesin-1 in the positioning of mitochondria (Wu et al., 2024) and spectrin (Glomb et al., 2023) in the DA9 neuron and kinesin light chain AID has validated the involvement of kinesin-1 in the outward movement of the spindle in meiosis I (Aquino et al., 2025).

Here, we characterise the effect of UNC-116 degradation on cargo transport by analysing DCV motility in the ALA neuron and on worm physiology by monitoring worm locomotion. UNC-116 protein levels drop after 1-3 hours of auxin treatment, and bidirectional DCV movement is inhibited over a similar timescale. Moreover, DCV transport in both directions recovers when worms are removed from auxin. Kinesin-1 therefore works as a crucial regulator of bidirectional DCV trafficking that, if lost, promptly halts transport. In contrast, worm locomotion is affected later, after 6-14 hours of degradation. UNC-116 loss has distinct effects on crawling and swimming over time, dependent on developmental stage, highlighting that the system can be used to provide insights about kinesin-1 involvement in different neuromuscular processes. Importantly, degrading UNC-116 specifically in neurons is sufficient to cause locomotion defects. Overall, we demonstrate that conditional depletion of UNC-116 can be used to dissect its fundamental functions in cargo transport and physiology on a variety of timescales.

## Results

### UNC-116 degradation is rapid and efficient

To establish if acute AID-mediated degradation of UNC-116 was feasible we used immunoblotting to monitor protein levels. An UNC-116 auxin-inducible degron strain was generated by inserting the minimal AID* tag (Zhang et al., 2015) followed by an ALFA tag (Götzke et al., 2019) at the 5’-end of the endogenous *unc-116* gene. The AID*-tagged strain, referred to here as *unc-116(deg)*, was crossed with a strain expressing a single copy of the TIR1 F-box gene downstream of the ubiquitous promoter *eft-3* (Ashley et al., 2021) to give *unc-116(deg);uTIR1*. When grown in the presence of auxin, TIR1 associates with the AID* and leads to the ubiquitination and proteasomal degradation of the target the protein complex in all cells (fig. 1A). We also crossed *unc-116(deg)* with a worm expressing TIR1 downstream of the pan-neuronal *rab-3* promoter *(nTIR1)* in an extrachromosomal array, as neuronal degradation of UNC-116 may be a better model for phenotypes caused by the loss of function of the neuron-specific human isoforms of kinesin-1 heavy chain, *KIF5A* and *KIF5C* (de Ligt et al., 2012; Dutta et al., 2018; Ebbing et al., 2008; Liu et al., 2014; Poirier et al., 2013).

**Figure 1.**
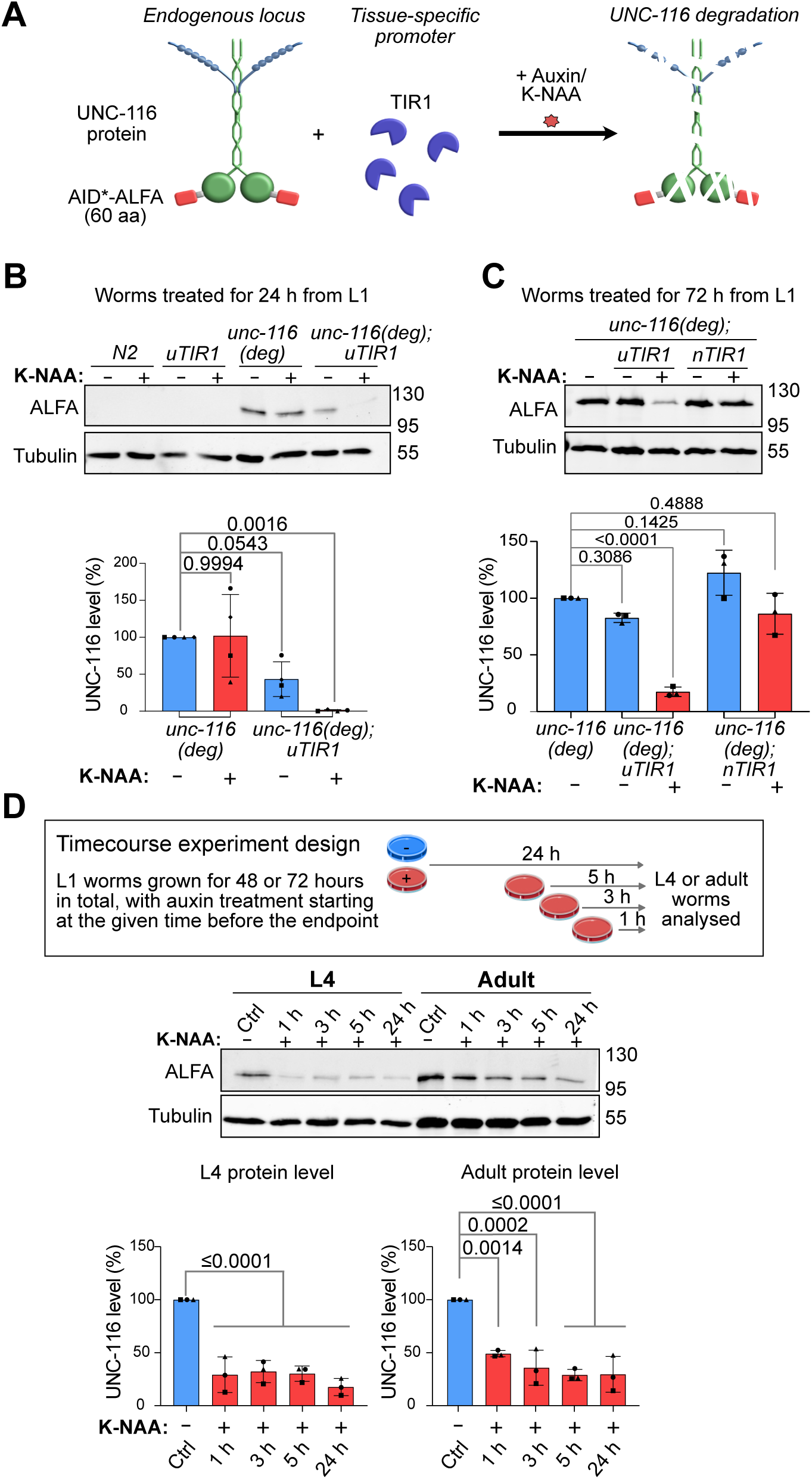
Robust UNC-116 degradation is seen upon K-NAA treatment. (A) Schematic of UNC-116 degradation using the AID system. UNC-116 is shown in green and kinesin light chains in blue. (B, C) Synchronised L1 worms of the indicated strains were grown ± K-NAA for 24 hours (B) or 72 hours (C). Immunoblotting was used to assess UNC-116 degradation using anti-ALFA antibodies with anti-tubulin as a loading control. Quantification was normalised to *unc-116 (deg) (-)* and anti-tubulin. (D) Time course of UNC-116 protein degradation in L4 larvae and adult *unc-116(deg);uTIR1* worms grown with K-NAA for the indicated times compared with untreated worms. Means ±SD are shown (N=4 for B, N=3 for C and D) with P-values from one-way ANOVA followed by post-hoc Dunnett test.

The UNC-116 protein was undetectable in *unc-116(deg);uTIR1* following treatment of L1 larvae with synthetic auxin (K-NAA) for 24 hours (fig. 1B) and greatly reduced when L1 worms developed to adults in K-NAA for 72 hours (fig. 1C). A decrease in UNC-116 protein was also seen following neuronal-specific degradation for 72 hours in *unc-116(deg);nTIR1* worms (fig. 1C). Since AID systems can demonstrate basal degradation by TIR1 in the absence of auxin (Hills-Muckey et al., 2022; Kanke et al., 2011; Natsume et al., 2016; Negishi et al., 2022; Schiksnis et al., 2020), we compared UNC-116 levels in the absence of K-NAA in the *unc-116(deg)* strains with and without uTIR1 expression (fig. 1B-C). While the presence of uTIR1 reduced UNC-116 levels somewhat in young larvae without K-NAA, the difference was not statistically significant.

Next, we investigated the rate of ubiquitous UNC-116 degradation in L4 and adult worms. To do this, L1 worms were allowed to develop for 48 or 72 hours, with K-NAA treatments starting 1, 3, 5 or 24 hours before the end point (fig. 1D). UNC-116 degradation was detected within 1 hour, and appeared more rapid in L4 worms, suggesting that speed of degradation is influenced by life stage, as reported in Zhang et al. (2015). The small residual pool of UNC-116 that remained after 24 hours of K-NAA treatment may correspond to the germline pool of UNC-116, which has important meiotic functions (Ellefson & McNally, 2009; McNally et al., 2012; Yang et al., 2005). The *eft-3* promoter driving TIR1 expression has been previously reported to express at low levels in the germline, similarly leading to a residual pool after degradation of the dynein heavy chain (Zhang et al., 2015).

An open question in neurobiology is whether phenotypes of neurodegeneration can be reversed if a functional transport machinery is restored, or whether transport loss leads to irreversible downstream defects (e.g. d’Ydewalle et al., 2011; Zhu & Sheng, 2011). Therefore, we investigated whether UNC-116 protein levels recovered after auxin removal. Worms were grown on K-NAA plates for 24 hours, followed by recovery on fresh NGM plates for different periods of time (fig. 2A). Some rescue of UNC-116 protein level was detected, but this was statistically insignificant compared to worms treated with K-NAA for 24 hours (fig. 2B). Since the rate of recovery of AID-tagged protein could be influenced by the duration of auxin exposure, adult worms were left on K-NAA plates for only 3 hours, and the recovery experiment was repeated. Protein level recovery was better after this shorter original K-NAA exposure, but still incomplete (fig. 2C).

**Figure 2.**
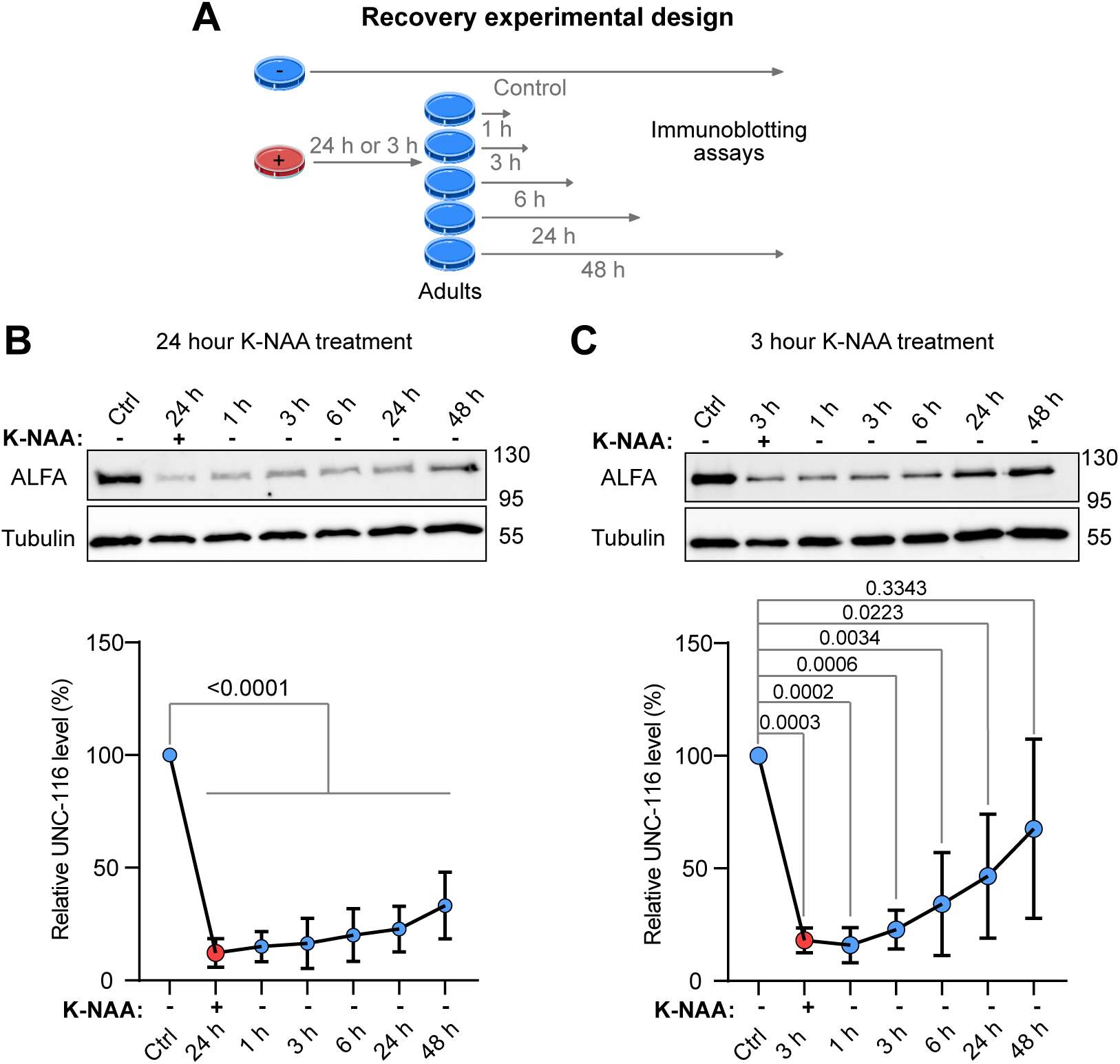
Recovery of UNC-116 protein levels after removal of K-NAA. (A) Time course of UNC-116 protein recovery in *unc-116(deg);uTIR1* adults treated with K-NAA for 24 hours (B) or 3 hours (C) followed by rescue for the time indicated, assessed using anti-ALFA antibodies. Quantification of ALFA-AID*-UNC-116 protein level is shown, normalised relative to anti-tubulin and the control ALFA signal (ctrl), which was from adults age-matched to the 48-hour time point. Means ±SD (N=3 for B, N=4 for C) and P-values from one-way ANOVA followed by post-hoc Dunnett test are shown.

Altogether, these results demonstrate that UNC-116 protein is efficiently and rapidly degraded in the presence of auxin in larval and adult worms, with statistically insignificant levels of background degradation without auxin. UNC-116 protein levels could recover following auxin treatment, but the extent was influenced by the duration of original auxin exposure.

### Bidirectional DCV transport is rapidly reduced by UNC-116 degradation and recovers after a rescue period

We have previously demonstrated that DCV movement in both directions is profoundly inhibited in the ALA interneuron in the hypomorphic *unc-116(rh24sb79)* mutant (Gavrilova et al., 2024) making DCV transport an excellent direct indicator of kinesin-1 function. To test the effect of acute UNC-116 loss on DCV movement using AID, we crossed the *unc-116(deg), unc-116(deg);uTIR1* and *unc-116(deg);nTIR1* strains with the *ida-1::gfp* strain, which expresses the GFP-tagged DCV transmembrane protein IDA-1 in the ALA, VC, HSN and PHC neurons (Cai et al., 2004; Zahn et al., 2001, 2004). ALA is ideal for imaging as has its cell body in the head and two laterally symmetrical axons running the length of each side of the body, with a further short neuronal process in the head (fig. 3A) (Sanders et al., 2013).

**Figure 3.**
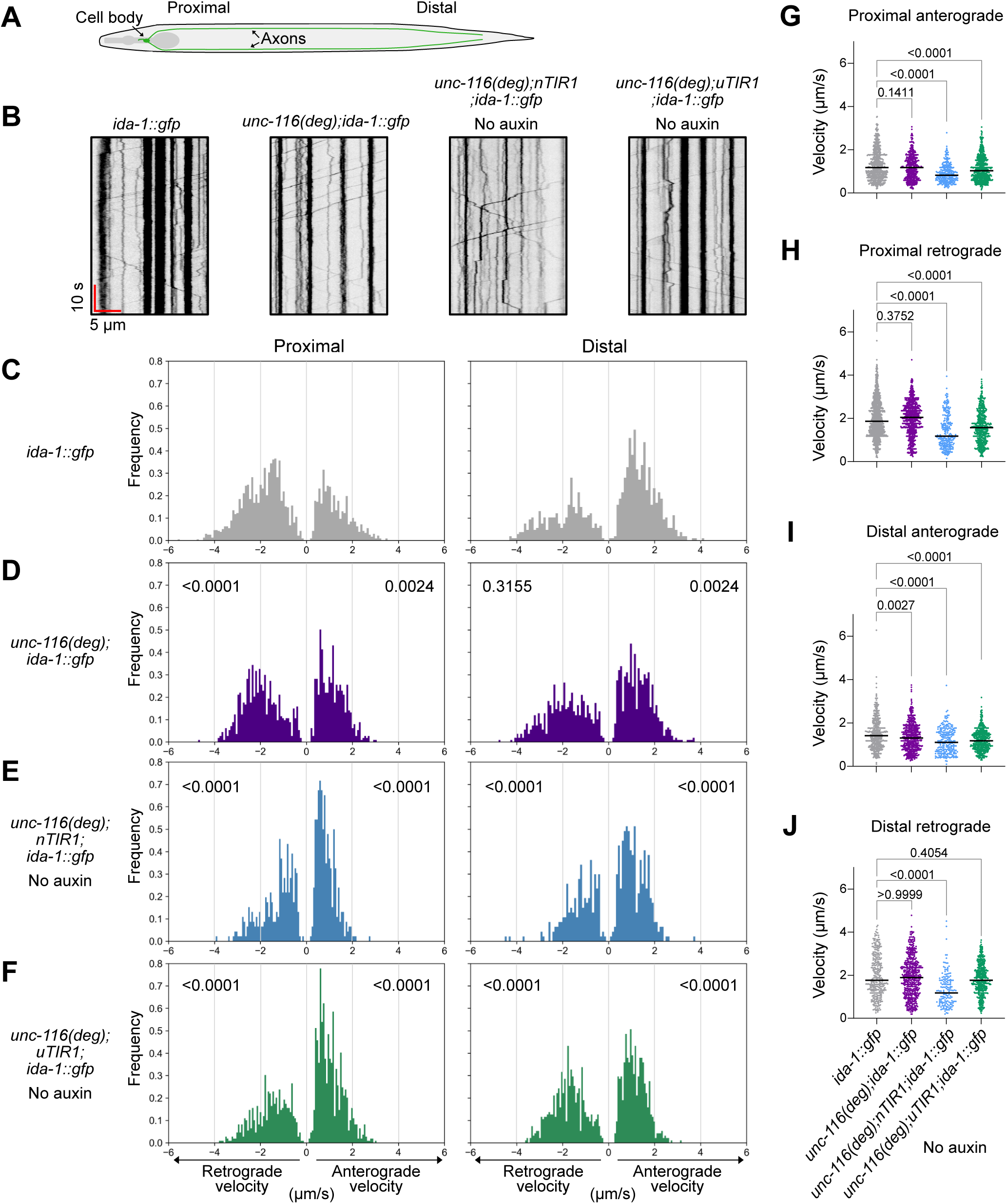
Effect of UNC-116 degron tagging and background degradation on DCV transport. (A) Schematic of the ALA neuron. (B) Examples of kymographs of DCVs in the proximal ALA neuron with time and distance scales indicated. The head is to the left. (C-F) Distributions of moving DCV track segment velocities in day 1 adults of *ida-1::gfp* (C: grey), *unc-116(deg);ida-1::gfp* (D: purple), *unc-116(deg);nTIR1;ida-1::gfp* (E: cyan), and *unc-116(deg);uTIR1;ida-1::gfp* (F: green) strains grown without K-NAA. The Y-axis displays the probability density function (frequency) with the X-axis showing segment velocity (negative values being retrograde, positive values being anterograde). The number of worms, kymographs and segments used to generate velocities are given in table S1. P-values from two-sample K-S tests are shown for movement in both directions, compared to the equivalent data in *ida-1::gfp*. (G-J) The same segment velocities plotted in scatter plots, displaying the median. Strains for all panels are indicated on the X-axis in J, with colour coding the same as in (C-F). P-values from Kruskal-Wallis tests followed by Dunn’s post-hoc test are shown, comparing each group to *ida-1::gfp*.

Spinning disc confocal recordings of DCV movement were taken in both the proximal and distal regions of the axon. As previously, DCVs were observed as either motile or stationary (Gavrilova et al., 2024; Goodwin et al., 2012; Ramirez-Suarez et al., 2019; Zahn et al., 2004). Axons in *C. elegans* have a plus-end-out microtubule polarity (Harterink et al., 2018), allowing assessment of anterograde and retrograde movement from kymographs (fig. 3B). DCV tracks were automatically identified using KymoButler (Jakobs et al., 2019) and then divided into anterograde, retrograde, and stationary segments (table S1, fig. S1A) using our Python analysis pipeline (Gavrilova et al., 2024). The velocities of the moving segments were plotted as velocity distributions (fig. 3C-F) and compared to *ida-1::gfp* using two-sample Kolmogorov-Smirnov (K-S) tests. In addition, Kruskal-Wallis tests were carried out to compare the difference in the rank sums between the different groups, used as an indicator of differences in median velocities (Ostertagová et al., 2014) (fig. 3G-J).

The presence of the AID*-ALFA tag on UNC-116 had no obvious effect on the proportion of moving DCVs (fig. S1A), although there was a slight shift towards lower anterograde velocities in *unc-116(deg);ida-1::gfp* compared to *ida-1::gfp* worms and the proportion of fast retrograde velocities in the proximal axon increased somewhat (fig. 3D). The only median velocity affected was for distal anterograde transport (fig. 3I). The AID*-ALFA tag therefore had little effect on UNC-116 function. Since background degradation reduced UNC-116 levels somewhat (fig. 1), we also analysed DCV transport in the AID*-UNC-116 TIR1 strains in the absence of auxin. Active DCV transport was seen in both the *unc-116(deg);uTIR1;ida-1::gfp* and *unc-116(deg);nTIR1;ida-1::gpf* strains (fig. 3, fig. S1A), although the number and percentage of moving DCVs were slightly lower in *unc-116(deg);nTIR1;ida-1::gfp* worms (table S1, fig. S1A), and fewer DCVs moved at the highest velocities (fig. 3C, 3E, 3F). Most median velocities were correspondingly slightly reduced (fig. 3G-J). Taken together, tagging UNC-116 with an AID*-ALFA tag and the low level of background degradation by TIR1 in the absence of auxin caused only minor changes in motility.

In stark contrast, UNC-116 degradation in the presence of auxin rapidly and drastically inhibited DCV motility in *unc-116(deg);nTIR1;ida-1::gfp* and *unc-116(deg);uTIR1;ida-1::gfp* worms (fig. 4, fig. S2). Already after 1 hour of degradation, the number of both anterograde and retrograde movements was greatly reduced in both degron strains, and further decreased after 4 and 24 hours, when moving DCVs were rare (fig. 4A, 4B, fig. S1B, S2A, S2B, movie 1). Note that segment counts, rather than frequencies, are plotted in fig. 4B-C and S2A because the number of movements differs greatly between conditions. A few anterograde movements persisted, even after 24 hours of K-NAA treatment, whereas retrograde motility was almost completely lost. Interestingly, after 1 hour of degradation the median anterograde velocities significantly increased in both the proximal and distal axon in *unc-116(deg);nTIR1;ida-1::gfp* worms, and in the distal region of *unc-116(deg);uTIR1;ida-1::gfp* worms (fig. 4D, 4F, fig. S2F, table S2, table S3). This may suggest that anterograde transport by kinesin-3/UNC-104 was initially maintained after UNC-116 degradation, after which this faster transport was also greatly reduced. In contrast, retrograde velocities were only significantly reduced in the proximal axon in *unc-116(deg);nTIR1;ida-1::gfp* worms after 1 hour K-NAA treatment (fig. 4E, 4G). Too few retrograde segments were identified at 4 and 24 hours to allow statistical comparison of the velocity distributions by K-S test with untreated DCV motility.

**Figure 4.**
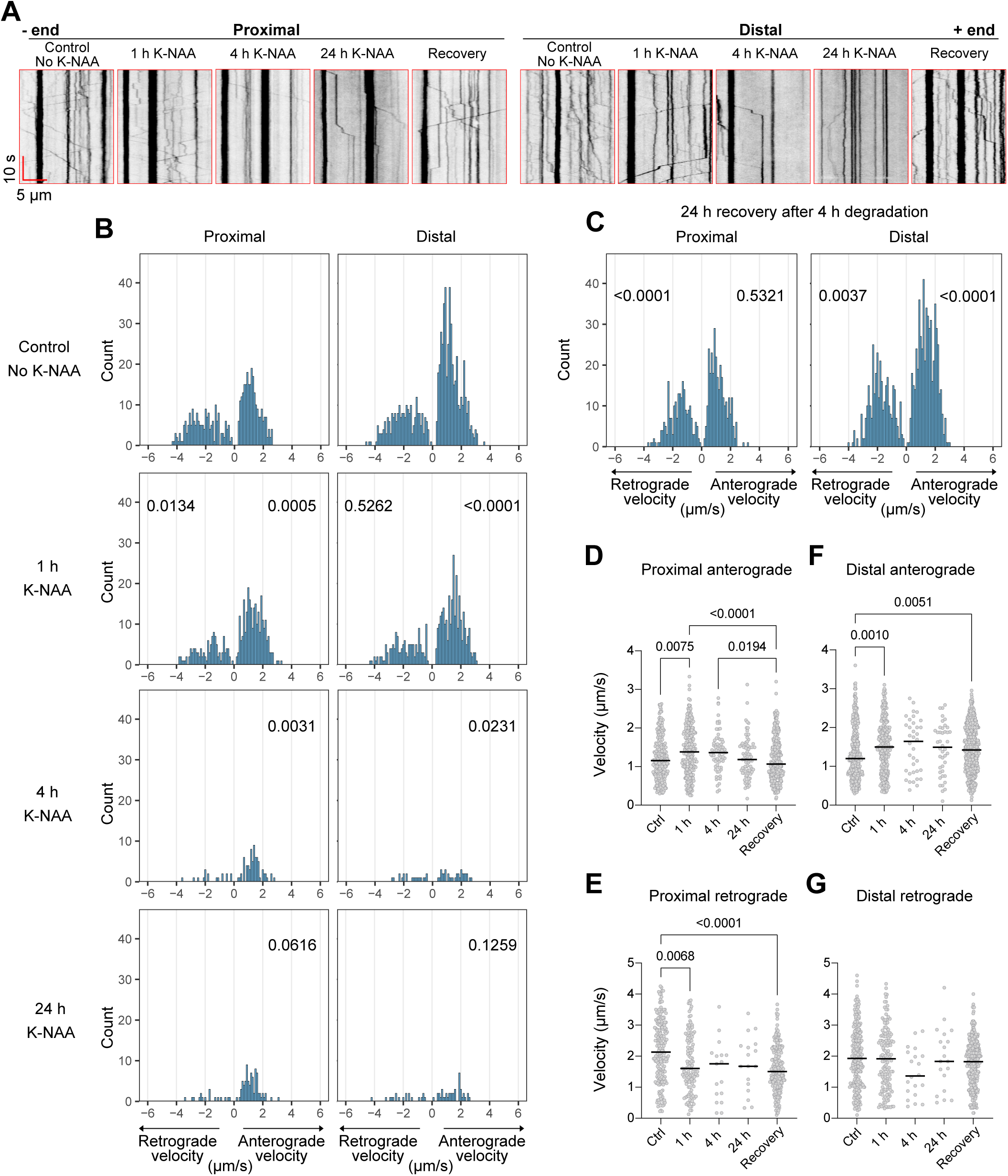
Both anterograde and retrograde DCV transport rapidly decline after neuronal UNC-116 degradation and recover upon K-NAA removal. (A) Example kymographs showing DCV movement along the proximal or distal region of the ALA neuron in *unc-116(deg);nTIR1;ida-1::gfp* worms under the indicated conditions. The head is to the left. (B) Velocity distributions of moving DCV track segments after incubation without (control) or plus K-NAA for 1, 4 or 24 hours with the number of moving segments (count) on the Y-axis. Number of worms, kymographs, and segments used to generate velocities are given in table S2. P-values are shown from two-sample K-S tests comparing retrograde and anterograde velocities in each condition and location to the equivalent subset in the untreated worms. The statistical analysis excluded the retrograde data from 4 h and 24 h, due to the limited number of values. (C) Velocity distributions of moving DCV track segments in worms treated with K-NAA for 4 h, then moved to K-NAA-free plates to recover for 24 h. P-values from two-sample K-S tests are shown, compared to the equivalent subset in the untreated day 1 adults. (D-G) The same data plotted in scatter plots displaying P-values ≤0.05 from Kruskal-Wallis tests followed by Dunn’s post-hoc test.

Finally, we investigated if DCV transport recovered when *unc-116(deg);nTIR1;ida-1::gfp* worms were treated with K-NAA for 4 hours, then transferred to regular NGM plates (without auxin) for 24 hours. Indeed, the number of DCV movements in both directions increased strongly, with the best recovery seen in the distal region (fig. 4A, 4C; fig. S1B; table S2). Interestingly, the anterograde velocities returned to control levels in the proximal axon but were slightly faster than controls in the distal axon. In contrast, retrograde velocities also recovered fully in the distal region but were slower in the proximal region.

Overall, degradation of UNC-116 for as little as one hour notably affected DCV movement in both directions along the length of the ALA neuron, and movement was profoundly stalled after four hours. These changes were reversed when worms were rescued from K-NAA.

### Swimming and crawling ability is drastically reduced by UNC-116 degradation

Worms with hypomorphic *unc-116* mutations are uncoordinated (Patel et al., 1993; Yang et al., 2005). The hypomorphic *unc-116(rh24sb79)* mutant has markedly reduced ability to crawl on stiff agar and swim/thrash in liquid (Gavrilova et al., 2024). This could be due to direct effects on cargo transport in neurons or muscle cells, but the presence of the partially compromised motor in all cells from birth may affect microtubule organisation or have pleiotropic effects on neuronal development that affect their function. We show above that motility of the exemplar cargo, DCVs, is rapidly inhibited upon UNC-116 degradation. This led us to ask if loss of ubiquitous kinesin-1 function at different developmental stages leads to locomotion defects, and if so, over what timescale.

We compared swimming/thrashing ability and crawling ability of adults that were allowed to develop from L1 for 72 hours (to adulthood) on plates with or without K-NAA (movie 2, movie 3). The number of body bends per second (a measure of swimming) and average crawling speed of *unc-116(deg);uTIR1* were significantly reduced upon UNC-116 degradation, to the same extent as the *unc-116(rh24sb79)* mutant (fig. 5A, 5B). Importantly, there was no significant difference in the swimming and crawling abilities of *unc-116(deg)* worms compared to *N2,* indicating that the AID*-ALFA tag on UNC-116 protein did not affect locomotion. However, *unc-116(deg);uTIR1* worms raised on plates *without* K-NAA had reduced swimming ability compared to *N2*, but normal crawling speed (fig. 5A, 5B). Thus, the effects of basal UNC-116 degradation by TIR1 in the absence of K-NAA could be detected by significantly reduced swimming but not crawling ability.

**Figure 5.**
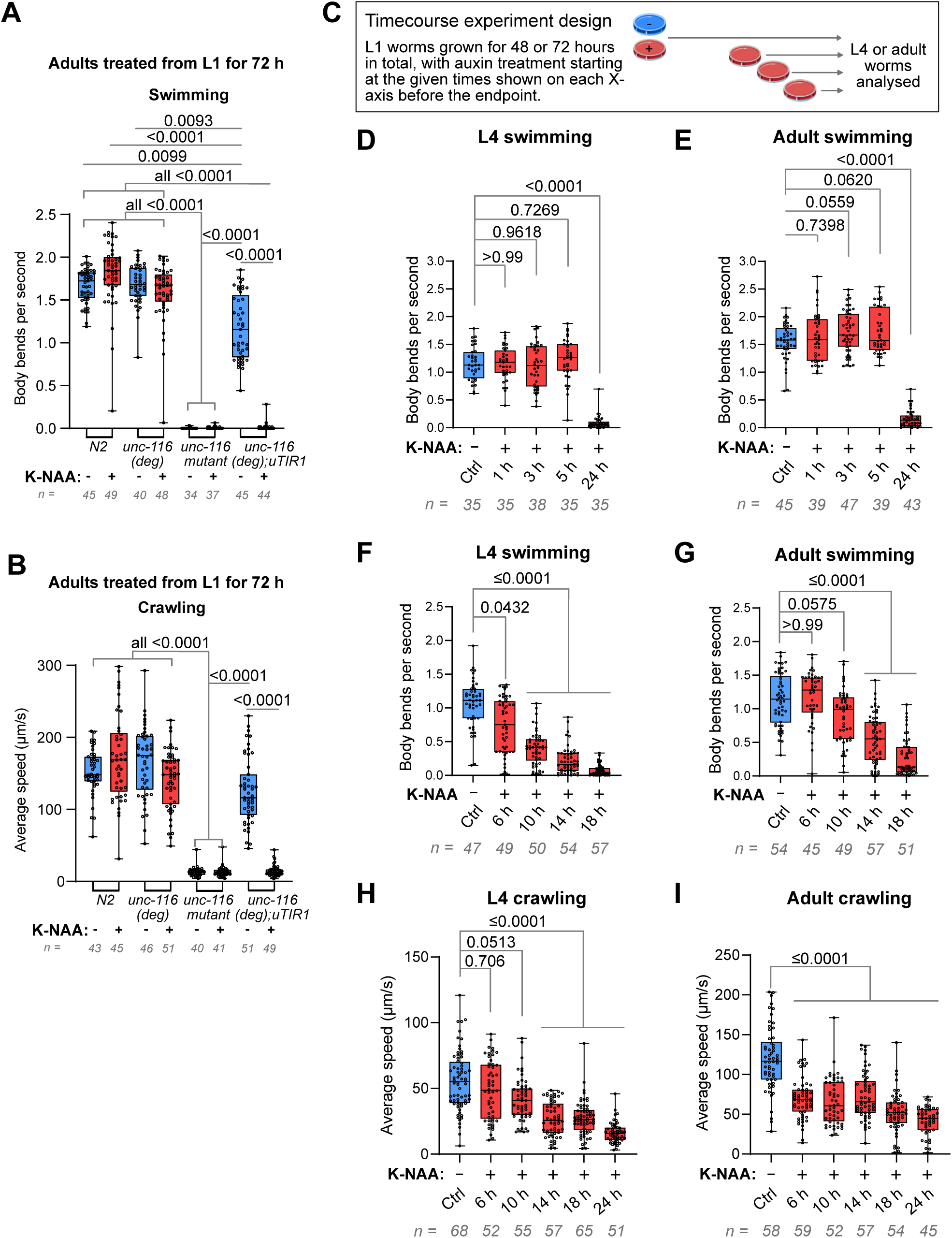
Effects of UNC-116 degradation on worm swimming and crawling. (A) Swimming ability (body bends per second) and (B) crawling ability (average speed) of adults grown ± K-NAA for 72 hours after L1 synchronization. (C) Experimental design of time course analysis of locomotion. Swimming ability measured as body bends per second, in L4s (D) or adults (E) using the same time points as for fig. 1D. Expanded time course analysis of swimming ability (F, G) and crawling speed (H, I), for L4 worms (F, H) and adults (G, I). All graphs show combined data from three biological repeats, with the total number of worms analysed indicated. Boxes display the median and IQR with whiskers from the minimum to the maximum value. Statistical analysis was by Kruskal-Wallis test followed by post-hoc Dunn’s test. Only P-values ≤0.05 are indicated on the graphs in (A) and (B).

We next asked how initiating UNC-116 degradation in L4 and adult worms affected locomotion, and if so, how long it took for the locomotion phenotypes to manifest. We performed time course studies of crawling and swimming in *unc-116(deg);uTIR1* L4 and adult worms treated with K-NAA for various times (fig. 5C). No significant change in swimming ability was seen after 1-5 hours of K-NAA treatment in adult and L4 worms (fig. 5D, 5E), even though UNC-116 protein was degraded (fig. 1D). After 24 hours of treatment, however, swimming ability was strongly reduced (fig. 5D, 5E). By assaying additional time points, we found that L4s showed a significant reduction in body bends per second after 6 hours of K-NAA treatment, while a significant reduction for adults was evident only after 14 hours of treatment (fig. 5F, 5G). When the effects on crawling were assessed, a somewhat different pattern was seen. A significant reduction in average crawling speed was observed already after 6 hours degradation in adults, but only after 14 hours degradation in L4s (fig. 5H, 5I). These data show that being able to degrade UNC-116 over time at specific developmental stages can reveal unexpected differences in sensitivity of behavioural traits to kinesin-1 loss.

Since DCV motility recovered when worms treated with auxin for 4 hours were transferred to auxin-free plates (fig. 4C), we wanted to test if locomotion would recover after auxin removal. However, because 5 hours of treatment had no effect on crawling and swimming (fig. 5D, 5E), we exposed *unc-116(deg);uTIR1* worms to K-NAA for 24 hours and then allowed them to recover without auxin for 1, 3, 6, 24 or 48 hours. Swimming ability did not recover (fig. 6A), whereas crawling velocity was slightly, but significantly, improved after 24 hours of recovery (fig. 6B). After 48 hours, crawling ability worsened again, perhaps due to aging effects.

**Figure 6.**
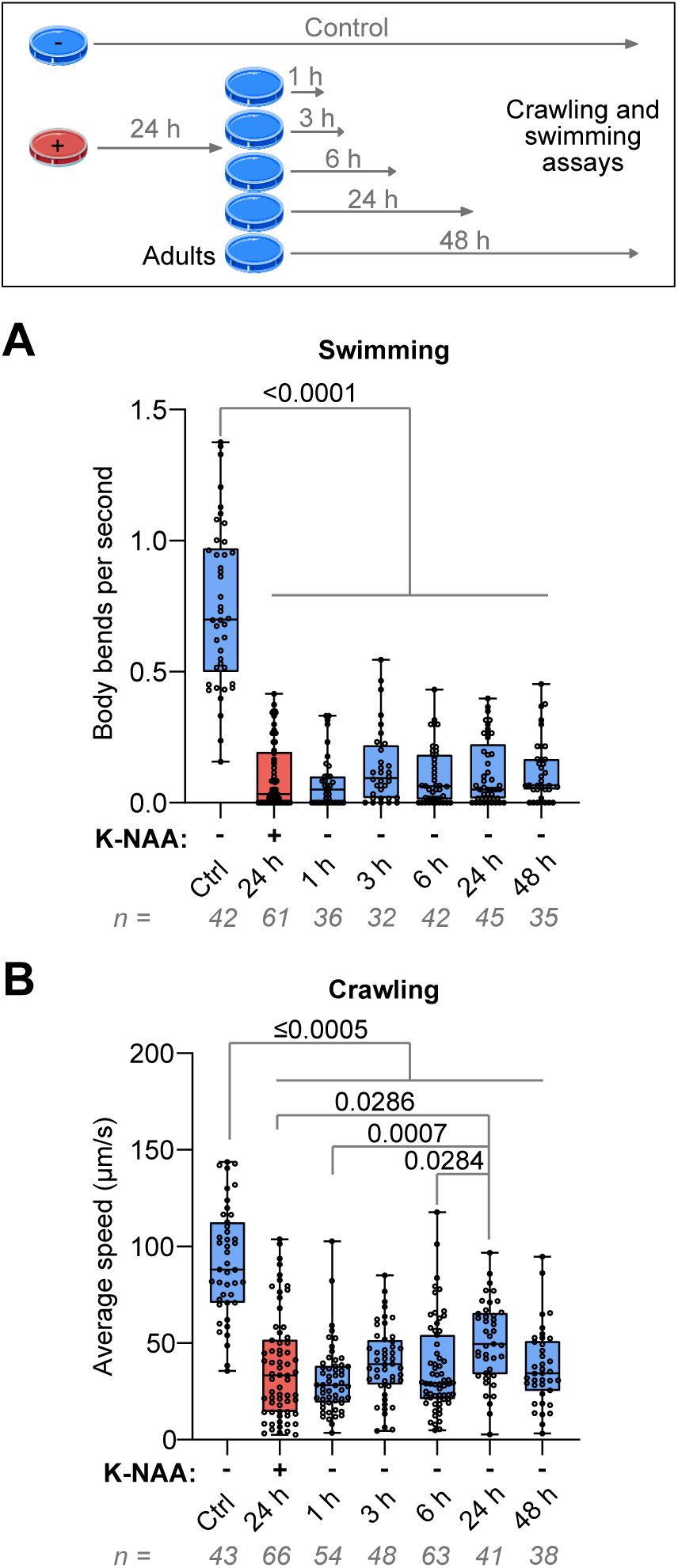
Limited recovery of locomotion after removal of K-NAA. Time course analysis of swimming (A) and crawling (B) in *unc-116(deg);uTIR1* adults treated with K-NAA for 24 hours followed by incubation without K-NAA for the time indicated. The number of worms analysed across three independent experiments is indicated, with boxes displaying the median and IQR with whiskers showing the data range. Statistical analysis was by Kruskal-Wallis test followed by Dunn’s post-hoc test. Only P-values ≤0.05 are indicated on the graphs.

The above experiments used worms expressing TIR1 in all cells. To test if degrading UNC-116 in neurons alone was sufficient to cause locomotion defects, we carried out swimming and crawling assays in adult *unc-116(deg);nTIR1* worms (fig. 7A, 7B, movie 4, movie 5). Strikingly, treatment with K-NAA for 24 hours (from the L4 stage to adulthood) resulted in markedly reduced swimming and crawling ability, similar to that seen in *unc-116(deg);uTIR1* worms. Interestingly, the effect of background degradation on swimming seen with ubiquitous TIR1 expression was not observed with neuronal TIR1 (fig. 7A). Together, these data indicate that UNC-116 function in neurons is necessary for normal locomotion.

**Figure 7.**
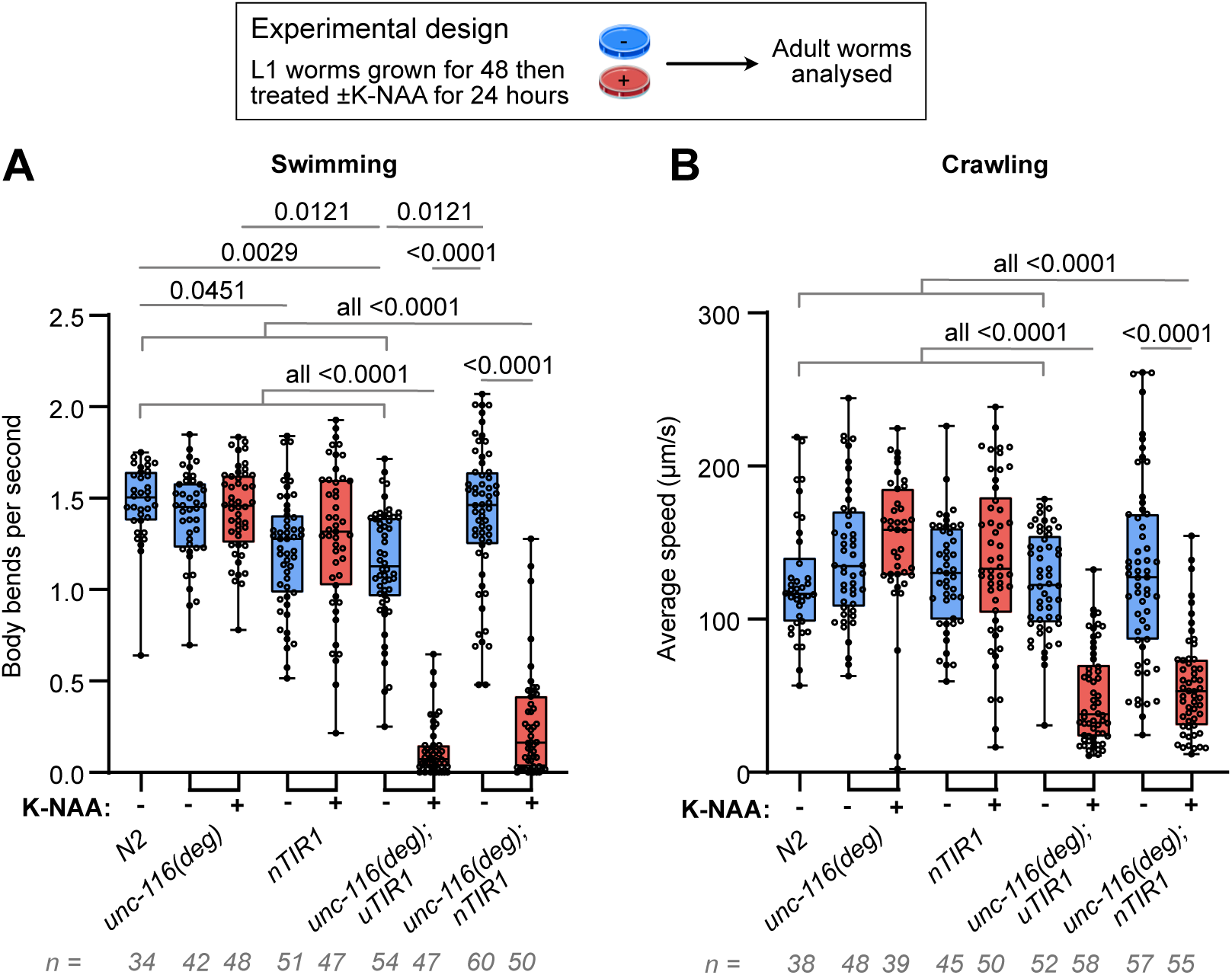
Neuronal UNC-116 degradation leads to swimming and crawling defects. Swimming ability (body bends per second) (A) or crawling ability (average speed) (B) of adult *unc-116(deg);nTIR1* worms grown on plates ± K-NAA for 24 hours. Graphs represent combined data from three biological repeats, with the number of worms shown. Boxes display the median and IQR with whiskers from the minimum to the maximum value. Only P-values ≤0.05 from the Kruskal-Wallis statistical test with Dunn’s post-hoc test are shown.

In summary, both ubiquitous and pan-neuronal UNC-116 degradation once development is complete leads to locomotion defects, but on a longer timescale than seen for loss of DCV motility. Furthermore, although DCV movement recovered well after removal of auxin, recovery of locomotion was limited after long-term UNC-116 degradation over the time course analysed.

## Discussion

AID offers exciting opportunities for investigating the function of essential proteins by controlled degradation. We have harnessed this to study the role of kinesin-1 in worms at selected developmental stages, exploring two levels of biological complexity: DCV motility and worm locomotion. We find that UNC-116 is degraded rapidly, and that bidirectional DCV movement is inhibited on a similar timescale. In contrast, it takes much longer for worm locomotion to be affected. Interestingly, the onset of crawling versus swimming defects was age dependent.

### Kinesin-1 and the regulation of dense core vesicle transport

This work shows conclusively that the dependence on kinesin-1 for efficient bidirectional transport of DCVs (Gavrilova et al., 2024) is independent of any developmental defects in the hypomorphic mutants used in that study. Thus, when the *C. elegans* nervous system has been allowed to develop normally, the post-developmental loss of the UNC-116 protein rapidly leads to the disruption of bidirectional DCV transport.

Different motors often colocalise on cargoes, sometimes binding via the same adaptor proteins that regulate unidirectional transport by selective motor activation (eg. Canty et al., 2023; Cason & Holzbaur, 2023; Fenton et al., 2021; Kendrick et al., 2019). Indeed, studies have reported the co-transport of opposing motors on DCVs. For example, in rat hippocampal neurons, the kinesin-3 family member KIF1A remains associated with DCVs during retrograde transport (Lo et al., 2011). Moreover, in *D. melanogaster* neurons kinesin-1 and kinesin-3 colocalise on DCVs (Lim et al., 2017), also seen in Rab6-positive secretory vesicles in mammalian cells, which are similar to neuronal DCVs (Serra-Marques et al., 2020). This indicates that all three opposing motors may be bound to DCVs at the same time. If this is the case, then why doesn’t the degradation of UNC-116 on DCVs lead to an initial increase in dynein-mediated transport of DCVs, before detrimental mislocalisation of anterograde-destined cargoes has had time to occur? Either earlier time points are needed to capture this, or other mechanisms can explain this behaviour.

One possibility is that a shared adaptor that binds both kinesin-1 and dynein is responsible for co-ordinating the direction of movement, but only when both motors are present. The specific adaptor(s) mediating this is unclear, although BICD is a candidate, as its isoforms have been found to associate with different motors on cargoes including DCVs and Rab6-positive vesicles (Ali et al., 2025; Grigoriev et al., 2007; Hoogenraad et al., 2001; Matanis et al., 2002; Schlager et al., 2010; Schlager et al., 2014). Other adaptors, such as JIPs, metaxins and TRAKs, may play the same role on other cargoes (Canty et al., 2023; Celestino et al., 2022; Zhao et al., 2021). However, since recent work has shown that dynein and its associated complex dynactin are transported anterogradely on different cargoes in cultured human i3 neurons (Fellows et al., 2024), it is also possible that a transport-ready dynein-dynactin complex is not present on kinesin-1-driven DCVs, which therefore cannot move retrogradely when UNC-116 is degraded.

Our results, and those we reported recently (Gavrilova et al., 2024), imply that kinesin-1/UNC-116 is the main anterograde motor for DCV transport in *C. elegans*, contrary to previous reports that kinesin-3/UNC-104 plays a critical role (Barkus et al., 2008; Lo et al., 2011; Zahn et al., 2004). Previous studies have highlighted the role of kinesin-3/UNC-104 in the delivery of DCVs out of the cell body and through the initial axon (Gumy et al., 2017; Lim et al., 2017; Park et al., 2023). Indeed, we find that a small proportion of anterograde DCV movement persists at later times, particularly in the proximal axon (fig. 4A, 4B; fig. S1A; fig. S2). Furthermore, after one hour, both proximal and distal anterograde velocities were significantly *increased* (fig. 4D, 4F), also visualised by the peak shift toward faster anterograde velocities in fig. 4B. This observation indicates that the faster motor UNC-104 (Kita et al., 2024) may be able to transport DCVs when some UNC-116 protein remains, but that robust UNC-116 degradation leads to an overall loss of transport. Curiously, in the recovery condition, anterograde velocity in the distal axon (fig. 4F) remained higher than the control. This might indicate that the fast motor kinesin-3/UNC-104 is active throughout the length of the axon in *C. elegans*.

Together, the results suggest that an intricate balance between motors is important, and potentially a mechanism by which they regulate each other. Previously, kinesin-1 but not kinesin-3 was found to engage in tug-of-war with dynein (Serra-Marques et al., 2020), and kinesin-3 detaches more easily from microtubules than kinesin-1 (Arpağ et al., 2014; Arpağ et al., 2019; Norris et al., 2014). Thus, association with microtubules by UNC-116 may be required for UNC-104 on DCVs to efficiently bind to microtubules and transport the vesicle, a model which has been previously suggested (Hancock, 2014). Importantly, because we are degrading tagged wild-type protein, rather than using partial loss-of-function mutants, we can rule out dominant-negative effects on bidirectional transport caused by an aberrant motor.

### The role of kinesin-1 in locomotion

Studies that assess phenotypes in adult *unc-116* mutants face the problem that kinesin-1 plays important roles in *C. elegans* development, meaning that it can be difficult to interpret the root cause of any defects seen. Neuronal development begins during *C. elegans* embryogenesis and continues at the L1 and L2 stages (Altun and Hall, 2011; Hedgecock et al., 1987; Sulston et al., 1983). By the L4 stage, the expression of genes involved in neurite development drops (Godini et al., 2022). UNC-116 is essential for organising dendritic microtubules during early development, which is crucial for subsequent trafficking of cargoes and organelles (Harterink et al., 2018; He et al., 2020; Yan et al., 2013). UNC-116 is also important for the transport of factors involved in axonal outgrowth (Drozd and Quinn, 2023; Lai and Garriga, 2004; Sakamoto et al., 2005; Su et al., 2006). Microtubule sliding by kinesin-1 is crucial for establishing initial neurite polarity and outgrowth, after which the microtubule cytoskeleton is largely immobilised (He et al., 2020; Lu et al., 2013).

By using AID, we have degraded UNC-116 in L4 and adulthood, when the nervous system is fully developed and axonal-dendritic microtubule polarity established, and find that this generates an uncoordinated phenotype. We also demonstrate that UNC-116 degradation in neurons alone of adult worms is sufficient to cause locomotion defects. This loss of mobility is likely due to failure in delivery of cargoes needed to maintain locomotory circuit function. Synaptic vesicle delivery should not be affected however, as they are transported by UNC-104/kinesin-3 (Hall and Hedgecock, 1991; Kumar et al., 2010; Niwa et al., 2016; Ou et al., 2010; Wu et al., 2013; Zheng et al., 2014).

Worm locomotion is principally controlled by cholinergic motoneurons that trigger muscle contraction on one side of the worm while GABAergic motoneurons cause simultaneous muscle relaxation on the opposite side (Cohen and Sanders, 2014; Haspel et al., 2020; Zhen and Samuel, 2015). This relies on inputs from a number of mechanosensory and command interneurons controlling forward and backward motion (Gjorgjieva et al., 2014). Glutamate receptor signalling is important in the command interneurons such as the AVA, which then activate cholinergic motor neurons (Haspel et al., 2020; Hoerndli et al., 2013; Hoerndli et al., 2015; Von Stetina et al., 2005). UNC-116 in a complex with KLC-2 transports vesicles containing AMPA-type glutamate receptor (AMPAR) subunits to postsynaptic sites in the AVA (Hoerndli et al., 2013; Hoerndli et al., 2015). Both *klc-2* mutants and worms with glutamate receptor mutations are hypersensitive to levamisole (Gavrilova et al., 2024; Sadananda & Subramaniam, 2021), a paralysing agent that activates one class of muscle acetylcholine receptor causing muscle hypercontraction. Thus, defective AMPAR transport in command interneurons that have lost UNC-116 may be one explanation for the gradual loss of locomotion. Kinesin function is also likely to be of direct importance in motoneurons, since AID of UNC-116 in the DA9 cholinergic motoneuron has been stated to lead to uncoordinated movement, although the phenotype was not analysed (Glomb et al., 2023).

Studies in mice have revealed that kinesin-1 is also involved in the transport of GABA_A_ receptors on vesicular cargo (Nakajima et al., 2012; Twelvetrees et al., 2010), as well as dopamine receptors (Cromberg et al., 2019). Furthermore, a number of neuropeptides influence worm locomotion, and worms with impaired DCV function are uncoordinated (Cai et al., 2004; Edwards et al., 2009; Flavell et al., 2013; Hu et al., 2011). Therefore, the consequent loss of DCV transport to release sites after UNC-116 degradation, as we observe in the ALA neuron, could contribute to the locomotion defects seen. Overall, loss of kinesin-1 function in several crucial neuromuscular pathways most likely leads to locomotion defects when UNC-116 is degraded after development. That the effects on swimming and crawling take much longer to manifest can be explained by kinesin-1 delivering cargoes such as receptors (which will turn over quite slowly) rather than synaptic vesicles, which must be constantly supplied to support neurotransmission directly. Future experiments where UNC-116 is degraded in specific subsets of neurons can further dissect the importance of kinesin-1 for locomotion.

### The timing of UNC-116 degradation-dependent effects on swimming and crawling alters with developmental stage

Unexpectedly, the onset of the loss of swimming ability after UNC-116 degradation occurred faster in L4s than in adults, whereas the loss of crawling ability occurred sooner in adults than L4s (fig. 5F-5I). Although evidence suggests that crawling and swimming are two extremes of one single gait (Berri et al., 2009; Boyle et al., 2012), the switch from crawling to swimming is influenced by serotonin signalling, whereas the switch from swimming to crawling is influenced by dopamine signalling (Pierce-Shimomura et al., 2008; Vidal-Gadea et al., 2011). Thus, the distinct effects of UNC-116 degradation on L4 and adult swimming and crawling could relate to differences in the availability of neurotransmitter and receptor reserves at neuromuscular junctions and synapses at different life stages.

Interestingly, RNA sequencing data from worms throughout the life cycle reveal that the expression of four out of seven serotonin receptors is higher in adults than in L4s (Boeck et al., 2016). This includes LGC-40 and SER-7, which are important for the crawl-to-swim transition and swimming velocity (Vidal-Gadea et al., 2011). Moreover, the expression of the serotonin reuptake transporter MOD-5 and the serotonin biosynthesis enzyme PAH-1 is higher in adults than in L4s. Conversely, the expression of three out of four dopamine receptors (DOP-1, DOP-2, DOP-3 but not DOP-4), the dopamine transporter DAT-1, and the enzyme controlling the rate-limiting step in dopamine biosynthesis (CAT-2) is decreased in adults compared to L4s (Boeck et al., 2016). This is consistent with the fact that serotonin promotes egg laying, whereas this is inhibited by dopamine signalling, so the balance between the two is important in adults (Weinshenker et al., 1995). If UNC-116 degradation disrupts trafficking pathways that subsequently affect the ability of serotonin and dopamine to regulate locomotion, deleterious effects on swimming ability will take longer in adults than L4s because the serotonin signalling machinery is expressed at higher levels, supporting local reserves. On the other hand, the onset of crawling defects will occur later in L4s because of higher expression of the dopamine signalling machinery. This provides a potential mechanistic explanation for the differential loss of crawling and swimming ability in L4s and adults, although further work is needed to test this hypothesis.

### Pros and cons of the UNC-116-AID system

The most obvious benefit of the UNC-116 AID system is that it avoids the use of mutants that have developed aberrantly and allows kinesin-1 to be removed from worms at any stage of development. In combination with tissue-specific expression of TIR1, kinesin-1 can be degraded only in specific cell types.

Another advantage is its suitability for studying the ability of worms to recover phenotypically after UNC-116 degradation. This could potentially provide information about whether disease symptoms caused by kinesin-1 impairment could be rescued by recovering the intracellular transport machinery. We found that the rate of UNC-116 protein recovery after degradation is influenced by the duration of auxin treatment (fig. 2B, 2C). Similarly, in Zhang et al. (2015), recovery rate was found to be dependent on auxin concentration. Worms treated with auxin for an extended period of time or with a high auxin concentration most likely have residual auxin left in tissues that takes a long time to wash out.

Consistent with this, we demonstrated that bidirectional DCV transport recovered when worms were removed from auxin plates after four hours of neuronal degradation and allowed to recover for 24 hours (fig. 4C). However, when assessing locomotion phenotypes after 24 hours of ubiquitous degradation, crawling ability recovered somewhat when worms were rescued from auxin for 24 hours, whereas swimming ability did not (fig. 6A, 6B). This may indicate that significant UNC-116 loss leads to certain irreversible locomotion defects. Alternatively, as mentioned, high residual auxin in tissues may delay phenotypic rescue.

Although the current UNC-116 AID system is a powerful tool, a limitation is the leaky degradation seen in the *unc-116(deg);uTIR1* strain in the absence of added auxin. Basal degradation of AID-fused proteins by TIR1 is proposed to be due to non-specific binding of TIR1 to the AID protein, or to the production of auxin-like indoles by commensal bacteria that can mimic auxin and trigger the association between TIR1 and the AID tag (Hills-Muckey et al., 2022; Kanke et al., 2011; Natsume et al., 2016; Negishi et al., 2022; Schiksnis et al., 2020). In *unc-116(deg);uTIR1,* basal degradation was detected at the protein level, slightly affecting swimming ability (fig. 1B, 1C; fig. 5A). The velocity of DCV transport was also reduced slightly by basal degradation (fig. 3D, 3E). Recent studies have developed improved variants of TIR1 for use in *C. elegans* by introducing a point mutation, TIR1^F79G^, that both improves the specificity of ligand binding and functions at very low concentrations of synthetic 5’Ph-auxin (Hills-Muckey et al., 2022; Negishi et al., 2022). Thus, the specificity of UNC-116 degradation can be improved in the future by crossing the *unc-116(deg)* strain with strains expressing TIR1^F79G^ in different tissues.

We found that the AID*-ALFA tag subtly affected DCV transport velocity (fig. 3D), emphasizing the importance of including the tagged motor on its own as a control. It also highlights the power of the DCV motility assay to detect very slight differences in motor activity. In contrast, UNC-116 degradation caused a rapid and dramatic loss of DCV motility. We have therefore demonstrated that conditional degradation of a microtubule motor can be used to study the functions of this motor in depth, including dissecting unknown aspects of cargo transport and systemic effects of motor loss. Extending this work to include AID of dynein and UNC-104 will allow rapid manipulation of each DCV motor, providing a testbed for unravelling how these three motors work together, without resorting to the use of mutants with partially functional motors.

## Materials and Methods

### *C. elegans* culture, strains, and genetics

*C. elegans* strains were cultured on nematode growth media (NGM) plates seeded with the *Escherichia coli* strain OP50 at 20°C, as previously described (Stiernagle, 2006). All strains used in this study are listed in Table S4. The *ida-1::gfp* strain was generated by Tobias Zahn in John Hutton’s laboratory, University of Colorado Health Sciences Center, Denver, USA, and was kindly supplied by Howard Davidson. The *unc-116(rh24sb79)* strain (Yang et al., 2005) was obtained from Frank McNally at the University of California-Davis, USA. The ubiquitous TIR1 strain (JDW225) (Ashley et al., 2021) was obtained from the *Caenorhabditis* Genetics Centre (CGC). The N2 Bristol variety was used as a wild-type reference strain. Strains generated in this study will be made available via the CGC.

The *unc-116(deg)* strain was generated by InVivo Biosystems using CRISPR-mediated insertion. Two sgRNAs targeting the 5’ end of the *unc-116* open reading frame were used to guide CRISPR/Cas9 (sgRNA1: AAAATGGAGCCGCGGACAGA + PAM and sgRNA2: CGGACAGACGGAGCAGAATG + PAM). A single-stranded oligodeoxynucleotide (ssODN) donor homology repair template was generated, containing the DNA sequence encoding AID*-ALFA and flanking homology arms (fig. S3). The sgRNAs were complexed with Cas9 protein prior to injection, followed by microinjection of the complete CRISPR/Cas9 gene editing mix into the gonads of young adult hermaphrodites. F1 animals were screened for the presence of co-CRISPR phenotype (*dpy-10 (cn64)*, Chromosome II). Homozygotes were confirmed by PCR and Sanger sequencing and backcrossed three times to N2.

The *nTIR1* extrachromosomal array strain was generated by InVivo Biosystems by microinjection of a mix of 15 ng/μl pNU3232 (*Prab-3::3xFLAG::TIR1::tbb-2 3’UTR*), 2 ng/μl pNU3225 (*Pmyo-2::NLS::GFP::unc-54 3’UTR*), 15 ng/μl pNU936 (Punc-119::unc-119::unc-119 3’UTR) and 69 ng/μl salmon testes DNA into *unc-119 (ed3)* null worms, followed by selection of worms with GFP expression and phenotypic rescue.

### Genotyping by single-worm PCR

DNA was amplified using OneTaq Quick-load 2X Master Mix (M0486L, New England Biolabs) according to the manufacturer’s protocol. Edit primer pairs (5’-CAGGATCTACATCTGGATCACC-3’, 5’-GAGAGAATTGACACACCTGC-3’) were used to verify that worms contained the AID*-ALFA CRISPR insert. Flanking primer pairs (5’-ATTGCAGGCATTGTAAGGAGAAGC-3’, 5’-CATGTGAGAGAATTGACACACCTGC-3’) were used during crosses to confirm homozygotes for the CRISPR edit.

### Synchronisation by bleaching

Plates with high densities of gravid adult hermaphrodites were washed with M9 buffer (22 mM KH_2_PO_4_, 86 mM NaCl, 42 mM Na_2_HPO_4_, 1 mM MgSO_4_) followed by aspiration into 15 ml tubes. Worms were pelleted by centrifugation at 500 x g for 1 minute, followed by aspiration of most of the supernatant. 1 ml of bleaching solution (0.7 M NaOH and 10% sodium hypochlorite solution (v/v) in ddH_2_O) was added to the pellet and vortexed periodically for 5 minutes, ensuring disintegration of the mothers but not the eggs. Three M9 washes were carried out, pelleting the released eggs by centrifugation between each wash. After the final wash, 5 ml of M9 solution was added, and tubes were left at room temperature on a rocker for at least 24 hours, ensuring hatching and arrest at the L1 stage. For experiments at the L2 stage, L1 worms were pelleted, aspirated and allowed to grow on plates for 24 hours. For experiments at the L4 stage, L1s were grown for 48 hours, and for the adult stage, L1s were grown for 72 hours.

### Auxin treatment

Potassium 1-Naphthaleneacetate (K-NAA) (N0006, TCI chemicals) was diluted in ddH_2_O and kept as a 100 mM stock at 4°C. 1 mM K-NAA plates were prepared as previously described (Martinez and Matus, 2020). Because auxin inhibits the growth of OP50 (Zhang et al., 2015), a 50 μl drop of 10X concentrated OP50 was added to each plate and allowed to dry at room temperature on the day of use. Synchronised worms were added to K-NAA plates for variable durations depending on the experimental procedure.

### Generation of worm lysates and western blotting

Worms were transferred to fresh NGM plates without OP50. 100 L2s/condition, 30 L4s/condition, or 20 adults/condition were picked except in experiments with 72 h degradation (fig. 1C), where 25 auxin-treated *unc-116(deg);uTIR1* and *unc-116(deg);nTIR1* adults were transferred due to their smaller size, in order to obtain approximately equal amount of protein in each sample. Worms were washed off with M9 and added to Eppendorf tubes, followed by centrifugation at 500 x g for 1 minute and aspiration of the supernatant, leaving ∼20 µl with the worm pellet. 20 µl of 2X sample buffer was added to the tubes, followed by boiling at 95°C for 10 minutes and SDS-PAGE. The gel was transferred onto a 0.45 µm pore PVDF membrane (IPFL00010, Millipore) using a BioRad wet transfer system. Antibodies were made up in TBS-T (20 mM Tris/HCl, pH 7.6-7.8, 150 mM NaCl, 0.1% Tween-20) according to the dilutions in table S5. Washed membranes were visualised using a LI-COR Odyssey Fc scanner. Band intensities were quantified using the Fiji Gel Analyzer function.

### Crawling and swimming assays

Movies of crawling and swimming/thrashing behaviour were captured using a Leica M165 FC Fluorescent Stereo Microscope with an optiMOS Scientific CMOS Camera and PLANAPO 1.0x objective. Assays were based on protocols from Bayrakli et al., (2015) and Hahm et al. (2015). For crawling assays, 10-20 worms were placed on a fresh NGM plate without OP50. For swimming assays, worms were added, followed by the addition of M9 buffer. Image streams consisting of 500 frames with ∼16.6 fps exposure were captured using Micro-Manager. Movies were analysed using the Fiji plugin wrMTrck (Nussbaum-Krammer et al., 2015) to quantify the number of body bends per second and average crawling speed, calculated as the sum of the length of all movement vectors for each track (μm) divided by the time recorded (s). Each measurement represents one worm. Three independent repeats were performed for each assay, and the data from multiple movies was combined for each strain/condition. During time course analyses, time points were staggered and analysed within ∼1 hour of each other.

### Timelapse imaging and kymograph analysis

For the time course experiment, time points were staggered and imaged within 3-4 hours of each other in day 1 adults. The recovery condition was imaged a day later. To mount worms for imaging, 2% agarose pads were generated by placing a drop of melted agarose on a vinyl record followed by flattening by a glass slide, as outlined in Rivera Gomez & Schvarzstein (2018). Worms were immobilised in 10 mM tetramisole hydrochloride (T1512, Sigma-Aldrich) diluted in M9 and placed in the resulting grooves in the solidified agarose, ensuring that they were parallel to the image view, and then covered with an 18 mm coverslip. The edges of the coverslip were sealed with VALAP (a 1:1:1 mixture of vaseline, lanolin and paraffin).

2D timelapse images of dense core vesicle were acquired using a CSU-X1 spinning disc confocal (Yokagowa) on a Zeiss Axio-Observer Z1 inverted microscope with a 100x/1.30 Plan-Apochromat objective, Prime 95B Scientific CMOS (1200 x1200 11 µm pixels; backlit; 16-bit) camera (Photometrics) and motorised XYZ stage (ASI). The 488 nm laser was controlled using an AOTF through the laserstack (Intelligent Imaging Innovations (3i)). Slidebook 2023 (3i) was used to capture continuously streamed images of a single Z-plane for 500 frames with a frame rate of 156 or 184 ms. Proximal movies were captured up to 300 µm from the cell body and distal movies up to 300 µm from the axon tip.

Raw TIFF files were stabilized using the image stabilizer plugin for Fiji using the best suitable initial reference slice. Standard parameters for the image stabilizer were used. A segmented line was then superimposed over the axon, and kymographs were generated using the multi kymograph plugin. Tracks were identified using KymoButler (Jakobs et al., 2019). Subsequent track analysis and the generation of direction and velocity data were carried out as previously described (Gavrilova et al., 2024). The supplemental movies were exported as AVIs in ImageJ, then compressed and converted to MP4 format using the H.264 encoder in HandBrake (version 1.9.2, handbrake.fr) with contrast quality set at 22.

### Statistical analysis

DCV velocity distributions were compared to control distributions by the pairwise two-sample Kolmogorov-Smirnov (K-S) test in every case with ≥50 data points. The null hypothesis that the two distributions come from the same population was rejected at P>0.05. Velocities were also compared using a Kruskal-Wallis test followed by a Dunn’s multiple comparisons test, as not all groups were normally distributed.

Statistical analysis for immunoblots was performed using one-way ANOVA tests followed by a post-hoc Dunnett test. For the crawling and swimming assays, normality was assessed for each sample group using a Shapiro-Wilk normality test. In all assays, some groups were not normally distributed. Therefore, statistical analysis was performed using a non-parametric Kruskal-Wallis test followed by a Dunn’s multiple comparisons test.

## Acknowledgements

We are grateful to Howard Davidson (University of Colorado Health Sciences Center, Denver), Frank McNally (University of California, Davis) and the CGC for providing worm strains. The CGC is funded by the NIH Office of Research Infrastructure Programs (P40 OD010440). The 3i spinning disk microscope used in this study was purchased by the University of Manchester Strategic Fund. Special thanks go to Peter March for his help with the confocal imaging. Finally, we are grateful to Jamie Herzig for help with coding and Martin Lowe for comments on the manuscript.

## Data availability

Source data and new strains will be made available upon publication.

