## Supplementary material for "Auxin-inducible degradation of UNC-116 in *C. elegans* inhibits bidirectional dense core vesicle transport and worm locomotion on different timescales"

#### Movies

**Movie 1: DCV transport in the proximal ALA axon in *unc-116(deg);nTIR1;ida-1::gfp* after periods of K-NAA treatment and recovery.** DCV transport is displayed in untreated (control) worms, worms treated with K-NAA for 1, 4 and 24 hours, and worms that have been allowed to recover from K-NAA for 24 hours after 4 hours of treatment. Time (m:s) is indicated. Scale bar = 5 μm.

**Movie 2: Swimming ability is lost after UNC-116 degradation.** Binary masks of indicated worm strains grown ± K-NAA for 72 hours after L1 synchronization are displayed. Swimming/thrashing ability was analysed by the wrmTrck Fiji plugin, which assigns each worm with a number and counts number of body bends (b). Time (m:s) is indicated. Scale bar = 1 mm.

**Movie 3: Crawling ability is lost after UNC-116 degradation.** Binary masks of indicated worm strains grown ± K-NAA for 72 hours after L1 synchronization are displayed. Time (m:s) is indicated. Scale bar = 1 mm.

**Movie 4: Neuronal and somatic UNC-116 degradation leads to swimming defects.** Binary masks of day 1 adults of the indicated strains grown ± K-NAA for 24 hours. Time (m:s) is indicated. Scale bar = 1 mm; b=number of body bends.

**Movie 5: Neuronal and somatic UNC-116 degradation leads to crawling defects.** Binary masks of day 1 adults of the indicated strains grown ± K-NAA for 24 hours. Time (m:s) is indicated. Scale bar = 1 mm.

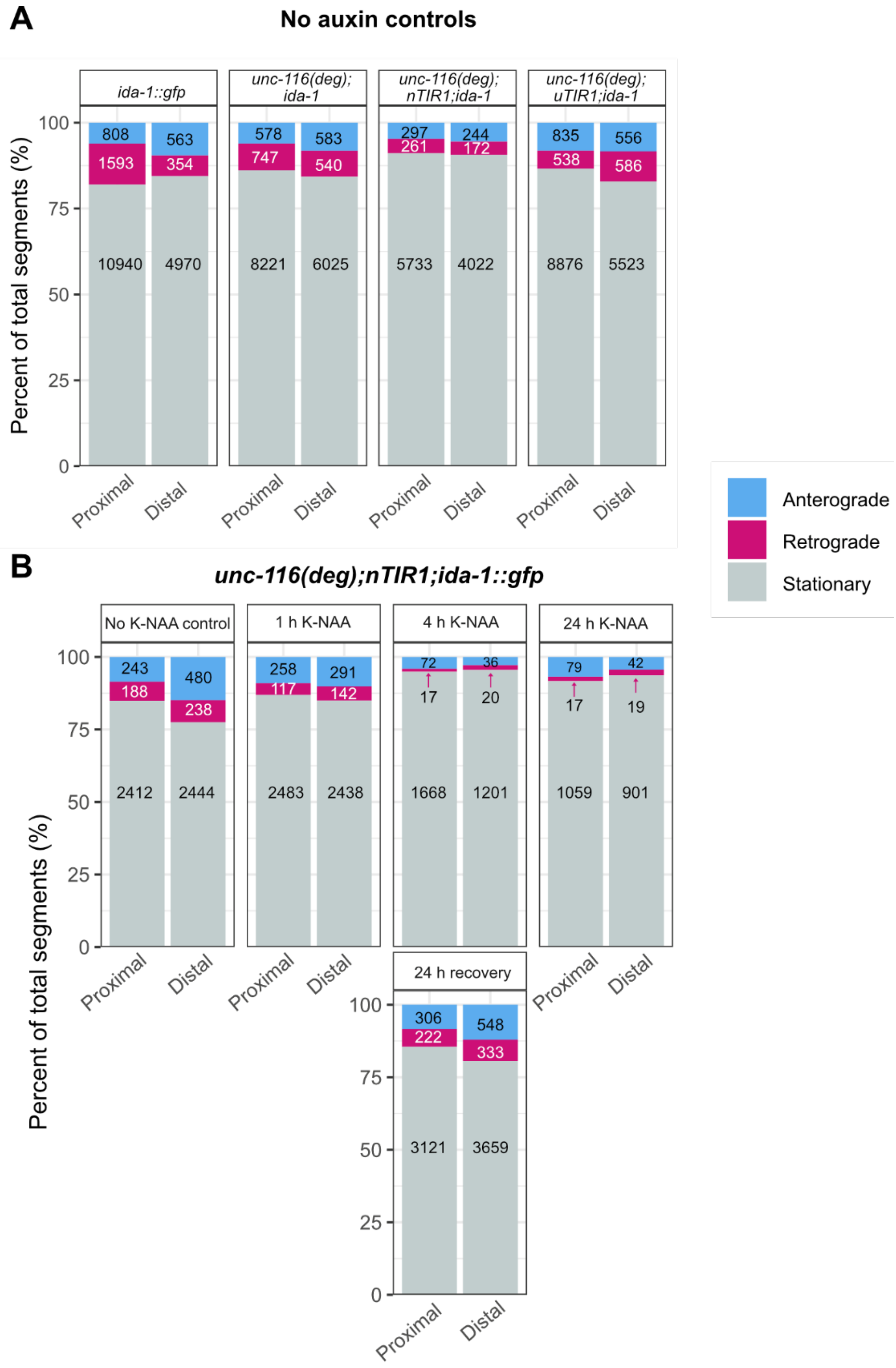


**Figure S1. Segment analysis from DCV tracks in the ALA axon identified from kymographs.** Proportion of anterograde, retrograde, and stationary segments identified in untreated strains (data from table S1) (A) and *unc-116(deg);nTIR1;ida-1::gfp* treated with K-NAA for indicated times (data from table S2) (B). Number of segments in each category is indicated on the graphs.

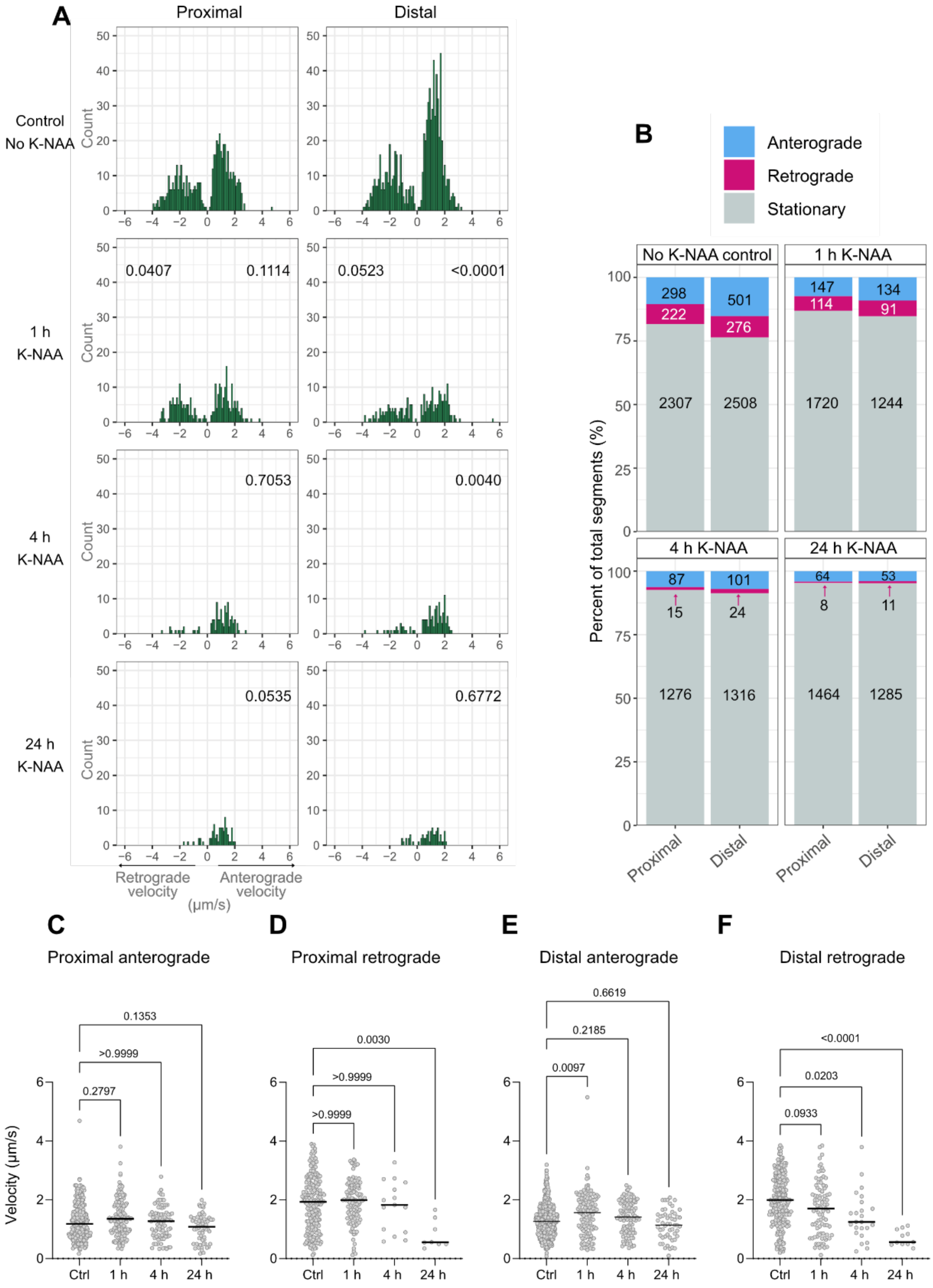


**Figure S2. Both anterograde and retrograde DCV transport rapidly decline after ubiquitous UNC-116 degradation.** (A) Velocity distributions of moving DCV track segments after incubation without (control) or plus K-NAA for 1, 4 or 24 hours with the number of moving segments (count) on the Y-axis. Number of worms, kymographs, and segments used to generate velocities are given in table S3. P-values are shown from two-sample K-S tests comparing retrograde and anterograde velocities in each condition and location to the equivalent subset in the untreated worms. The statistical analysis excluded the retrograde data from 4 h and 24 h, due to the limited number of values. (B) Proportion of anterograde, retrograde, and stationary segments identified in *unc-116(deg);uTIR1;ida-1::gfp* treated with K-NAA for indicated times (data from table S3). (C-F) The same data plotted in scatter plots displaying the P-values from Kruskal-Wallis tests followed by Dunn's post-hoc test are shown, comparing each group to the control group.

5’ -
GGTGATCATTATTCTCTGAAAATGGAGCCGCGGACAATGCCTAAAGATCCAGCCA
AACCTCCGGCCAAGGCACAAGTTGTGGGATGGCCACCGGTGAGATCATACCGG
AAGAACGTGATGGTTTCCTGCCAAAAATCAAGCGGTGGCCCGGAGGCGGCGGC
GTTCGTGAAGTCAGGATCTACATCTGGATCACCATCTAGACTCGAAGAGGAGCT
CCGCAGACGGCTTACCGAACCAGGGGGGACCGGATCCGGGTCCTCGACGTCG
ACGATGGAACCAAGAACTGATGGTGCTGAGTGCGGTGTCCAGGTATTAATTTTC
CCCGCCTAGAT - 3’

Homology arms, AID*, ALFA, Linkers, 27 bp recoded region with silent mutations to prevent re-cutting by sgRNAs

**Figure S3. Donor homology template for CRISPR insertion of AID*-ALFA tag at the *unc-116* locus.**

### Tables

**Table S1.** Summary of DCV motility analysis for strains without auxin treatment, from fig. 3.

| Condition | Number of worms | Number of kymographs | Number of tracks | Direction | Number of segments | Mean velocity (µm/s) | Median velocity (µm/s) | Min velocity (µm/s) | Max velocity (µm/s) |
|---|---|---|---|---|---|---|---|---|---|
| *ida-1::gfp* proximal | 17 | 90 | 6721 | Anterograde | 808 | 1.3 | 1.2 | 0.2 | 3.5 |
| | | | | Retrograde | 1593 | 2.0 | 1.9 | 0.2 | 5.6 |
| | | | | Stationary | 10940 | - | - | - | - |
| *ida-1::gfp* distal | 17 | 69 | 2967 | Anterograde | 563 | 1.5 | 1.4 | 0.1 | 6.3 |
| | | | | Retrograde | 354 | 2.0 | 1.8 | 4.3 | 0.3 |
| | | | | Stationary | 4970 | - | - | - | - |
| *unc-116(deg); ida-1::gfp* proximal | 19 | 70 | 4806 | Anterograde | 578 | 1.2 | 1.2 | 0.2 | 3.1 |
| | | | | Retrograde | 747 | 2.0 | 2.0 | 0.2 | 4.7 |
| | | | | Stationary | 8221 | - | | - | - |
| *unc-116(deg); ida-1::gfp* distal | 18 | 60 | 3613 | Anterograde | 583 | 1.3 | 1.3 | 0.3 | 3.7 |
| | | | | Retrograde | 540 | 1.9 | 1.9 | 0.2 | 4.8 |
| | | | | Stationary | 6025 | - | - | - | - |
| *unc-116(deg); nTIR1; ida-1::gfp* proximal | 17 | 70 | 3175 | Anterograde | 297 | 0.9 | 0.8 | 0.2 | 2.8 |
| | | | | Retrograde | 261 | 1.4 | 1.2 | 0.1 | 3.9 |
| | | | | Stationary | 5733 | - | - | - | - |
| *unc-116(deg); nTIR1; ida-1::gfp* distal | 17 | 52 | 2249 | Anterograde | 244 | 1.2 | 1.1 | 0.1 | 3.0 |
| | | | | Retrograde | 172 | 1.3 | 1.2 | 0.2 | 4.5 |
| | | | | Stationary | 4022 | - | - | - | - |
| *unc-116(deg); uTIR1; ida-1::gfp* proximal | 16 | 112 | 5177 | Anterograde | 835 | 1.1 | 1.0 | 0.1 | 3.0 |
| | | | | Retrograde | 538 | 1.6 | 1.6 | 0.2 | 3.8 |
| | | | | Stationary | 8876 | - | - | - | - |
| *unc-116(deg); uTIR1; ida-1::gfp* distal | 16 | 79 | 3390 | Anterograde | 556 | 1.2 | 1.2 | 0.3 | 3.2 |
| | | | | Retrograde | 586 | 1.8 | 1.8 | 0.2 | 3.6 |
| | | | | Stationary | 5523 | - | - | - | - |

**Table S2.** Summary of DCV motility analysis in the *unc-116(deg);nTIR1;ida-1::gfp* K-NAA timecourse experiment shown in fig. 4

| Condition | Number of worms | Number of kymographs | Number of tracks | Direction | Number of segments | Mean velocity (μm/s) | Median velocity (μm/s) | Min velocity (μm/s) | Max velocity (μm/s) |
|---|---|---|---|---|---|---|---|---|---|
| Control proximal | 12 | 20 | 1439 | Anterograde | 243 | 1.2 | 1.2 | 0.3 | 2.6 |
| | | | | Retrograde | 188 | 2.2 | 2.1 | 0.2 | 4.2 |
| | | | | Stationary | 2412 | - | | - | - |
| Control distal | 10 | 18 | 1614 | Anterograde | 470 | 1.3 | 1.2 | 0.3 | 3.6 |
| | | | | Retrograde | 238 | 2.0 | 1.9 | 0.1 | 4.6 |
| | | | | Stationary | 2444 | - | | - | - |
| 1 h proximal | 13 | 21 | 1437 | Anterograde | 258 | 1.4 | 1.4 | 0.3 | 3.3 |
| | | | | Retrograde | 117 | 1.8 | 1.6 | 0.1 | 3.8 |
| | | | | Stationary | 2483 | - | | - | - |
| 1 h distal | 12 | 21 | 1461 | Anterograde | 291 | 1.5 | 1.5 | 0.3 | 3.1 |
| | | | | Retrograde | 142 | 1.9 | 1.9 | 0.3 | 4.3 |
| | | | | Stationary | 2438 | - | | - | - |
| 4 h proximal | 13 | 19 | 885 | Anterograde | 72 | 1.4 | 1.4 | 0.4 | 2.8 |
| | | | | Retrograde | 17 | 1.5 | 1.8 | 0.2 | 3.6 |
| | | | | Stationary | 1668 | - | | - | - |
| 4 h distal | 13 | 19 | 634 | Anterograde | 36 | 1.5 | 1.6 | 0.4 | 2.8 |
| | | | | Retrograde | 20 | 1.5 | 1.4 | 0.4 | 2.8 |
| | | | | Stationary | 1201 | - | | - | - |
| 24 h proximal | 13 | 19 | 588 | Anterograde | 79 | 1.2 | 1.2 | 0.2 | 3.1 |
| | | | | Retrograde | 17 | 1.8 | 1.7 | 0.3 | 3.4 |
| | | | | Stationary | 1059 | - | | - | - |
| 24 h distal | 12 | 19 | 486 | Anterograde | 42 | 1.4 | 1.5 | 0.1 | 2.6 |
| | | | | Retrograde | 19 | 1.9 | 1.8 | 0.5 | 4.2 |
| | | | | Stationary | 901 | - | | - | - |
| Recovery proximal | 15 | 29 | 1850 | Anterograde | 306 | 1.2 | 1.1 | 0.2 | 3.2 |
| | | | | Retrograde | 222 | 1.6 | 1.5 | 0.1 | 3.7 |
| | | | | Stationary | 3121 | - | - | - | - |
| Recovery distal | 15 | 27 | 2310 | Anterograde | 548 | 1.4 | 1.4 | 0.1 | 3.0 |
| | | | | Retrograde | 333 | 1.8 | 1.8 | 0.2 | 4.0 |
| | | | | Stationary | 3659 | - | - | - | - |

**Table S3.** Summary of DCV motility analysis in the *unc-116(deg);uTIR1;ida-1::gfp* K-NAA timecourse experiment shown in fig. S2.

| Condition | Number of worms | Number of kymographs | Number of tracks | Direction | Number of segments | Mean velocity (µm/s) | Median velocity (µm/s) | Min velocity (µm/s) | Max velocity (µm/s) |
|---|---|---|---|---|---|---|---|---|---|
| Control proximal | 12 | 20 | 1426 | Anterograde | 298 | 1.3 | 1.2 | 0.2 | 4.7 |
| | | | | Retrograde | 222 | 1.9 | 1.9 | 0.1 | 3.9 |
| | | | | Stationary | 2307 | - | - | - | - |
| Control distal | 12 | 17 | 1617 | Anterograde | 501 | 1.3 | 1.3 | 0.1 | 3.2 |
| | | | | Retrograde | 276 | 2.0 | 2.0 | 0.2 | 3.9 |
| | | | | Stationary | 2508 | - | - | - | - |
| 1 h proximal | 11 | 17 | 996 | Anterograde | 147 | 1.4 | 1.4 | 0.3 | 3.8 |
| | | | | Retrograde | 114 | 1.9 | 2.0 | 0.1 | 3.4 |
| | | | | Stationary | 1720 | - | - | - | - |
| 1 h distal | 13 | 13 | 741 | Anterograde | 134 | 1.5 | 1.6 | 0.3 | 5.5 |
| | | | | Retrograde | 91 | 1.7 | 1.7 | 0.1 | 3.8 |
| | | | | Stationary | 1244 | - | - | - | - |
| 4 h proximal | 13 | 18 | 693 | Anterograde | 87 | 1.2 | 1.3 | 0.3 | 2.8 |
| | | | | Retrograde | 15 | 1.8 | 1.8 | 0.6 | 3.3 |
| | | | | Stationary | 1276 | - | - | - | - |
| 4 h distal | 13 | 15 | 725 | Anterograde | 101 | 1.4 | 1.4 | 0.4 | 2.5 |
| | | | | Retrograde | 24 | 1.4 | 1.2 | 0.2 | 3.8 |
| | | | | Stationary | 1316 | - | - | - | - |
| 24 h proximal | 12 | 19 | 771 | Anterograde | 64 | 1.1 | 1.1 | 0.2 | 2.0 |
| | | | | Retrograde | 8 | 0.8 | 0.6 | 0.3 | 1.7 |
| | | | | Stationary | 1464 | - | - | - | - |
| 24 h distal | 12 | 18 | 677 | Anterograde | 53 | 1.2 | 1.1 | 0.1 | 2.1 |
| | | | | Retrograde | 11 | 0.7 | 0.6 | 0.3 | 1.1 |
| | | | | Stationary | 1285 | - | - | - | - |

**Table S4.** *C. elegans* strains used in this study

| Strain name | Referred to as | Genotype | Source |
|---|---|---|---|
| Bristol N2 | *N2* | Wild-type | CGC |
| BL5752 | *ida-1::gfp* | *inIs182 (ida-1p::ida-1::gfp) I; inIs181 (ida-1p::ida-1::gfp) IV* | Zahn et al. (2004) |
| N/A | *unc-116 mutant* | *unc-116 (rh24sb79) III* | Yang et al. (2005) |
| OL0332 | *unc-116(deg)* | *unc-116(uk001[AID::ALFA::unc-116]) III* | This study |
| JDW225 | *uTIR1* | *wrdSi23 [eft-3p::TIR1::F2A::mTagBFP2::AID*::NLS::tbb-2 3'UTR] (I:-5.32)]* | CGC Ashley et al. (2021) |
| OL0334 | *nTIR1* | *ukEx31[rab-3p::3xFLAG::TIR1::tbb-2 + (pNU936) unc-119p::unc-119::unc-119u + (pNU3225) NLS::myo-2::GFP]; unc-119(ed3)* | This study |
| OL0388 | *unc-116(deg);uTIR1* | *unc-116(uk001[AID::ALFA::unc-116]) III); wrdSi23 [eft-3p::TIR1::F2A::mTagBFP2::AID*::NLS::tbb-2 3'UTR] (I:-5.32).* | This study |
| OL0389 | *unc-116(deg);nTIR1* | *unc-116(uk001[AID::ALFA::unc-116] III; ukEx31[rab-3p::3xFLAG::TIR1::tbb-2 + (pNU936) unc-119p::unc-119::unc-119u + (pNU3225) NLS::myo-2::GFP]; unc-119(ed3)* | This study |
| OL0390 | *unc-116(deg);uTIR1;ida-1::gfp* | *unc-116(uk001[AID::ALFA::unc-116] III); wrdSi23 [eft-3p::TIR1::F2A::mTagBFP2::AID*::NLS::tbb-2 3'UTR] (I:-5.32); inIs181; inIs182 [ida-1p::ida-1::gfp]* | This study |
| OL0391 | *unc-116(deg);nTIR1;ida-1::gfp* | *unc-116(uk001[AID::ALFA::unc-116] III; ukEx31[rab-3p::TIR1::tbb-2 + (pNU936) unc-119p::unc-119::unc-119u + (pNU3225) NLS::myo-2::GFP]; unc-119(ed3); inIs181; inIs182 [ida-1p::ida-1::gfp]* | This study |
| OL0479 | *unc-116(deg);ida-1::gfp* | *unc-116(uk001[AID::ALFA::unc-116] III); inIs182 (ida-1p::ida-1::gfp) I; inIs181 (ida-1p::ida-1::gfp) IV* | This study |

**Table S5.** Antibodies used

| Antibody | Host species | Dilution | Source | Product number |
|---|---|---|---|---|
| TAT1 (Tubulin) mAb | Mouse | 1:5000 | Keith Gull, University of Oxford | N/A |
| ALFA 800 CW | Camelid | 1:10,000 | NanoTag | N1502-Li800-L |
| Anti-Mouse 680 | Donkey | 1:10,000 | Jackson ImmunoResearch Laboratories | 715-625-151 |
